# Pharmacological impacts on schizophrenia functional analysis: a postmortem proteome study

**DOI:** 10.1101/2021.10.26.465949

**Authors:** Rawan S. Alnafisah, James Reigle, Sinead M O’Donovan, Adam J. Funk, Jaroslaw Meller, Robert E. Mccullumsmith, Rammohan Shukla

## Abstract

Schizophrenia (SCZ) is a severe and debilitating mental illness. Antipsychotic drugs (APDs) are used to treat both positive and negative SCZ symptoms, by influencing the cellular, subcellular-synaptic, and molecular processes. We posit that these effects influence our understanding of SCZ. To address this, we analyzed postmortem dorsolateral prefrontal cortex grey matter samples from control and SCZ subjects (n=10/group) using liquid-chromatography mass-spectrometry-based proteomics. We retrieved SCZ-altered and APD-influenced proteome-sets using linear and mixed linear models, respectively, and validated them experimentally using independent cohorts and insilico using published datasets. Functional analysis of proteome-sets was contrasted at the biological pathway, cell-type, subcellular-synaptic, and drug-target levels. The SCZ-altered proteome was conserved across several studies from DLPFC and other brain areas and was dependent on drug effect. At the pathway level, we observed an aberrant extracellular event and, except for homeostasis, signal-transduction, cytoskeleton, and dendrites associated downregulated changes, the APDs compensated for the majority of the SCZ-altered pathways. At the cell-type level, the up-and down-regulated SCZ-altered events were associated with two different subsets of striatum projecting layer-5 pyramidal-neurons regulating dopaminergic secretion. At the subcellular synaptic level, compensatory pre- and post-synaptic events were observed. At the drug target level, dopaminergic processes influence the SCZ-altered up-regulated proteome, whereas non-dopaminergic and a diverse array of non-neuromodulatory mechanisms influence the SCZ-altered down-regulated proteome. While these findings are dependent on pharmacological effects, they are also consistent with previous SCZ studies, implying the need to re-evaluate previous results. We discuss our findings in the context of cortico-striatal influence in SCZ-pathology.

## INTRODUCTION

Schizophrenia (SCZ) is a devastating mental disorder that typically emerges in late adolescence or early adulthood, and results in social and mental impairment^1^. SCZ-pathophysiology involves altered functionality of different brain areas. However, the dorsolateral prefrontal cortex (DLPFC), an area crucial for verbal and working memory, due to evidence of SCZ associated morphometric changes and neurotransmitter abnormalities^2–4^, has been of special interest. Direct examination of protein expression in the DLPFC of SCZ postmortem tissue using high throughput approaches has revealed several altered proteins and biological pathways. However, the main limitation of these high throughput approaches is the difficulty to regress the influence of antipsychotic-drugs (APDs)^5^, which influence cellular, subcellular, and molecular processes. Unless comparing drug naïve and drug treated SCZ subjects with control groups, it is hard to infer the true molecular correlates of SCZ.

To address this challenge, several studies, have performed parallel expression studies on non-human primates^6^ or rodents^7^, ^8^ that were chronically treated with either typical- or atypical-APD. While these animal-based studies identify APD-signatures (relevant gene or proteins influenced by APDs), they heavily rely on unsustainable causal assumption about SCZ-pathophysiology and/or the APDs mechanism of action (MOA). First, these animal-based studies focus on characterizing the SCZ symptoms rather than understanding the molecular bases of the disease. Second, healthy animals exposed to pharmacological treatment or lesions to mimic SCZ symptoms might induce changes in brain structure or composition that might not reflect the disease pathology. lastly, animal models metabolize APDs differently than humans, consequently, alter different metabolic pathways. Nevertheless, specific signatures derived from these models are important in pointing genes or proteins (henceforth: features) involved in APD action.

Recently, several resources have emerged that provide drug specific signatures. Particularly useful are the comparative toxicogenomics database (CTD)^9^ anchoring well curated drug-associated features, and connectivity-map (cmap)^10^, anchoring experimentally derived drug-specific features. We predict that these resources, together with known gene-ontologies and recent advances in generating cell-specific signatures, can give an opportunity to generate informed hypotheses regarding drug and disease effects at the cellular level in SCZ-pathology. To this end, we performed systematic integration of these resources with the liquid chromatography mass-spectrometry (LCMS) based proteomics data highlighting altered changes attributed to typical and atypical APD medications and SCZ pathophysiology in human postmortem DLPFC. We demonstrate unique proteins, biological-pathways, synaptic and cellular alterations influenced by the drugs and SCZ-pathophysiology, as well as potential MOAs connected with them.

## Material and methods

### Subjects and Tissue Preparation

Dorsolateral prefrontal cortex (DLPFC, Brodmann area 9) tissues from SCZ and non-psychiatrically control subjects were obtained from the Maryland Brain Collection and were distributed by the Maryland Brain Collection and the Alabama Brain Collection. There were two cohorts used in this study. A mass spectrometry cohort from SCZ (n=10) and non-psychiatrically ill controls (n=10) (Supplementary Table 1). Second, conformation study cohort of SCZ (n =23) and non-psychiatrically ill controls (n = 23) (Supplementary Table 1). Schizophrenia subjects were diagnosed based on DSM-IV criteria. The medical records of the subjects were examined using a formal blinded medical chart review instrument, and in person interviews with the subjects and/or their caregivers, as previously described^11^. 14μm thick tissue sections were generated and stored at −80 °C until used. Frozen tissues were thawed, scrapped, and homogenized in 1ml of 5mM Tris HCl, 32 M Sucrose pH 7.4, with 1% protease and phosphatase inhibitor (Halt, Thermofisher^™^). Protein concentration was measured with the Pierce BCA kit for mass spectrometry and western blot experiments detailed in the supplementary information.

### SCZ-altered and APD-influenced proteome

Protein intensity were quantile normalized using the *limma* package in R. To estimate the variance explained by each variable associated with the samples, we used *variancePartition,* an R package that uses a linear mixed model to summarize each variable’s contribution in terms of fraction of variance explained^12^. Unfortunately, the cohort size does not allow regressing the effect of all the variable^13–15^. Accordingly, only age and postmortem interval (PMI), which explained the highest variability across all the samples (Figure 1A) were regressed from the normalized data. This was implemented using a ~Age + PMI + phenotype as design in the *model.matrix* function of the *limma* package in R. Here, phenotype represents data from both control and SCZ samples. A p-value threshold of 0.05 was used to determine the SCZ-altered differentially expressed proteome (Supplementary Table 2).

**Figure 1:**
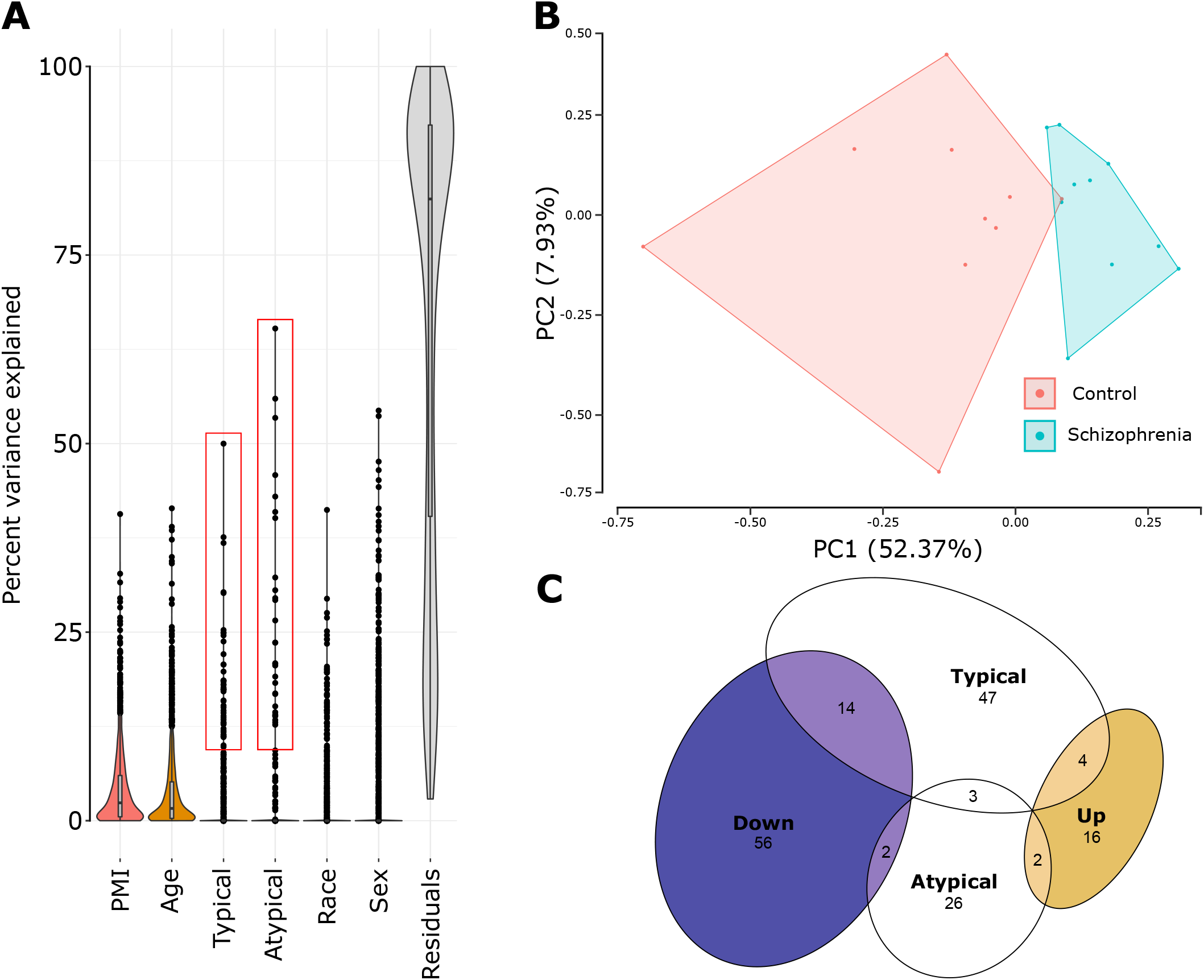
SCZ-altered and APD influenced proteome. **A)** A violin plot to sum up the proteome-wide trend and rank each variable’s contribution. Age and PMI explained the most percentage of variance in the data, followed by APDs, race, and sex. The influence of APDs was investigated using proteins that contributed more than 10% of the variables (red boxes). **B)** Principal components analysis segregating the proteome profile of the control and schizophrenia subjects. **C)** Venn Diagram showing intersection of up- and down-regulated SCZ-altered and APD-influenced proteome (red boxes in A). For more details on the proteome, see supplementary table 1.

For a given variable, the *variancePartition* fits a mixed-model to each protein across all sample, allowing per protein estimation of the explained variation across cohort. (Supplementary Table 3). We leveraged this to filter proteins explaining >10% variation associated with the two new variables constructed by classifying the administered APDs into “atypical” and “typical” class. The variable (“atypical” and “typical”) holds binary values with TRUE if the drug of that class was administered or FALSE if not. No subjects were treated with drugs of both the class. The filtered proteins for both the class were together called the APD-influenced proteome. Note that unlike the SCZ-altered proteome for which the directionality (up and down-regulated) is estimated based on the log fold change, APD-influenced proteome estimated based on percent variability explained, is directionless. Furthermore, as the APDs were delivered only to SCZ-subjects, the APD-influence are specific for SCZ-subjects.

### Theme-centric pathway analysis

The list of SCZ-altered, and APD-influenced proteins were employed to perform pathway analysis using gene ontology (GO) database. All pathway profiles (*q-value* < *0.05*) were tagged for their respective list identifier (i.e., up-SCZ, down-SCZ, atypical and typical) and stacked vertically in to one file. The enrichment score (ES) for each pathway was determine by −log_10_ transformation of the q-value (i.e., −log_10_(*q-value*)). To compare ES across SCZ-altered, and APD-influenced proteome, the vertically stacked pathway profile was pivoted using the *group.by* function in R (Supplementary Table 4). To summarize and filter the pathways associated with each list into neurobiologically relevant themes, we leveraged the hierarchical organization of the GO database and picked 19 GO terms (themes) belonging to biological process-, molecular function-, or cellular component-related ontologies representing most neurobiological functions of interest and then looked for the GO terms in our results, which are linked to these 19 themes as child nodes using *GOBPANCESTOR*, *GOMFANCESTOR*, and *GOCCANCESTOR* functions in R *GO.db* package. The ES of pathways associated with each list were used to make Figure 3.

### Brain area, cell-type, synapse, and drug-MOA/target specific enrichment analysis

While GO-based pathway analysis is useful for understanding functional changes at the biological-process, molecular-function, and cellular-component levels, it falls short of explaining central nervous system-specific functional alterations and druggable mechanisms. For this, we leveraged data available from several past and recent high-throughput studies focusing on similar proteomics analysis of the same (DLPFC)^16, 17^ and different brain areas^18–23^, synapse-^24^ and cell-specific^25^ changes, and drug-specific molecular alteration^9, 10^. Lists of features (genes or proteins) associated with these studies were curated (Supplementary Table 5) and their enrichment in the SCZ-altered, and APD-influenced proteome was determined using hypergeometric overlap analysis (i.e., fisher exact test) implemented by gene-overlap package in R-3.6.0. Given two gene-lists, a significance test for their overlap in comparison with a genomic background representing the universe of known genes (21,196 features, default used by the package) can be described using hypergeometric distribution. The null-hypothesis represents an odd-ratio <1 whereas the alternate hypothesis represents an odd-ratio >1.

Notably, the hypergeometric overlap analysis with the feature list of similar studies and drug can also be used to validate the SCZ-altered, and APD-influenced proteome. The drug-specific gene-sets were downloaded from drug signature database (DSigDB)^*26*^; http://dsigdb.tanlab.org/DSigDBv1.0/. DSigDB contains signatures from various sources, including those from cmap^*10*^, and CTD^*9*^. However, note that unlike cmap, which anchors drugs signatures with direction (i.e., up, or downregulated), the signatures form CTD are directionless. Due to the study’s objective of elucidating the MOA/targets linked with SCZ-altered proteome, only drugs with available MOA were considered. MOA for drugs were downloaded from https://clue.io/touchstone and http://ctdbase.org/downloads/.

## RESULTS

### Schizophrenia altered and APD-influenced DLPFC proteome

LCMS-based expression profiles for 1547 non-imputed proteins were obtained from postmortem DLPFC grey matter (layer 1 through 6) samples of control (CTL) and SCZ subjects (n=10/group; Supplementary Table 1). After regressing the effect of age and PMI, each accounting for >5% overall variability in the data (Figure 1A), we obtained 72 (*p-value* < 0.05) differentially expressed proteins (up= 16, down= 56, henceforth SCZ-altered proteome) which clearly segregate the control and SCZ samples (Figure 1B). The subjects in this study were treated either with typical or atypical APDs which together accounted for < 5% of overall variability in the data (Figure 1A). Since regressing this small effect from a limited number of samples is challenging^27, 28^, we instead filtered proteins which independently explained >10 % of variability associated with atypical and typical APDs using linear mixed modelling^12^ (Figure 1A, red boxes, see methods). Regardless of direction (i.e., up- or down-regulated), 26 and 47 proteins were associated with atypical and typical APDs, respectively (henceforth: APD-influenced proteome). Except for downregulated SCZ-altered and typical APDs there was minimal overlap between the SCZ-altered and APDs-influenced proteome (Figure 1C).

Next, we performed validation at multiple levels. First, the proteomics approach was validated using western blot analysis of glutaminase (*GLS*, Figure 2A), a protein differentially expressed in our MS based proteomics analysis (Figure 2B and Supplementary Table 2), in an independent cohort. There were no differences between CTL and SCZ (Figure 2C), however a significant downregulation was observed in the age range of 20 to 37 (Figure 2D). Second, the resulting up- and down-regulated differentially expressed proteins list were validated using hypergeometric overlap analysis with other similar proteomics study form DLPFC^16, 17^ (Figure 2F, top) and other brain areas including *insular-cortex^21^*, *hippocampus^22^*, *anterior-hippocampus^20^*, *genu-of-corpus-callosum^23^ and grey^18^ and white^19^ matter of anterior-cingulate-cortex (ACC)* (Figure 2B, middle). The upregulated proteins overlapped with studies from most other areas, while the downregulated proteins only overlapped with ACC grey matter but consistently overlapped with DLPFC in other studies. Notably, all studies overlapped significantly with features associated either with atypical or typical APDs. Lastly, we validated the atypical and typical APD-influenced proteome using hypergeometric overlap analysis with drug specific features available form either CTD or cmap (Figure 2B, bottom). Consistent with their mechanism involving dopaminergic transmission, typical APDs associated proteins overlapped significantly with *Haloperidol*— a dopamine receptor antagonist (and typical APD)^29^. Interestingly, *Haloperidol* also overlapped significantly with the upregulated proteins and were administered to the studied SCZ samples (Supplementary Table 1). Atypical APDs associated proteins overlapped with *valproic-acid^30^* and *clomipramine^31–33^* which are often used to treat SCZ. Lastly, the downregulated proteins overlapped with *alprazolam*, which has shown marked improvement in treating positive and negative schizophrenic symptoms^*34*^.

**Figure 2:**
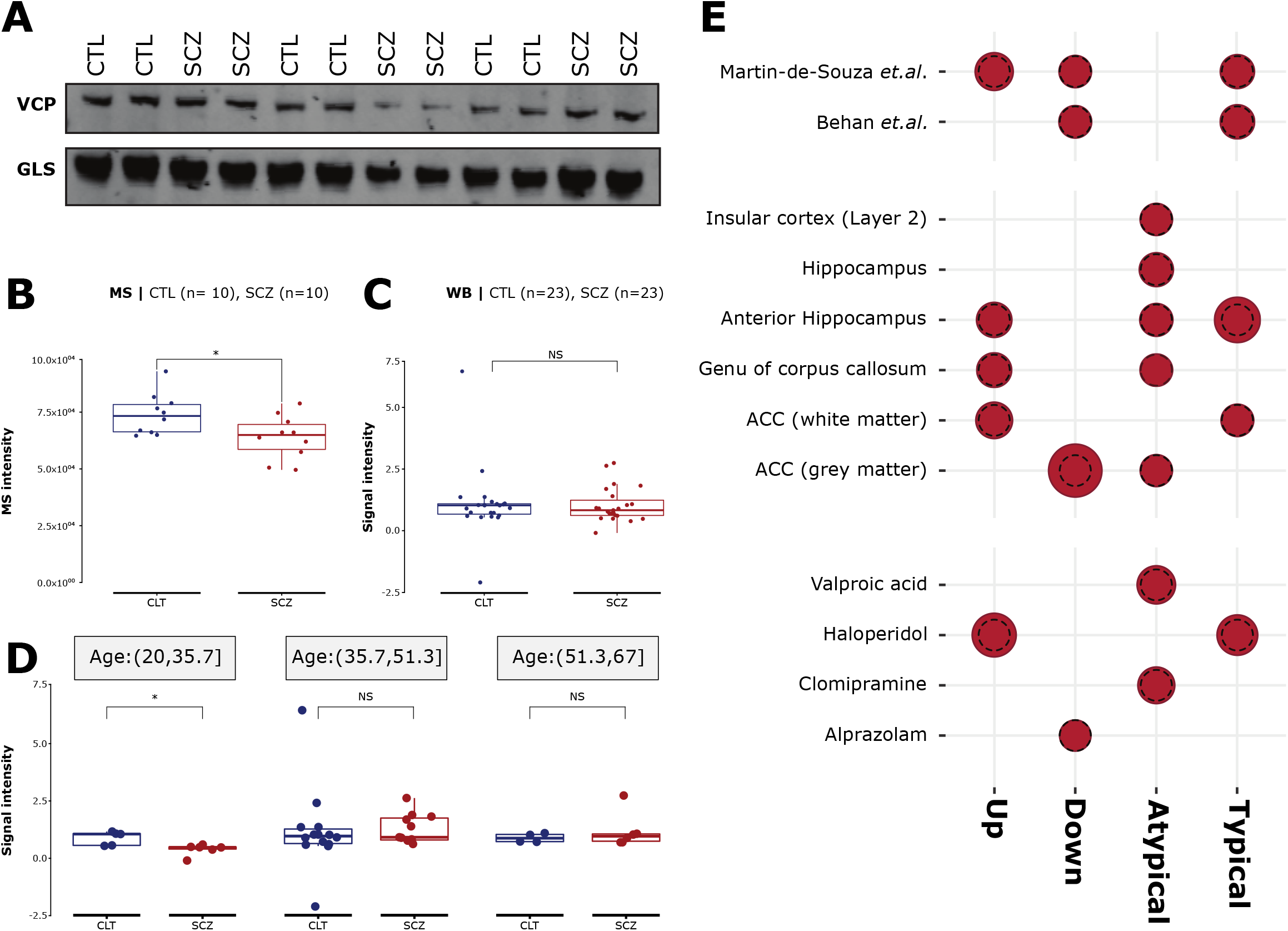
Validation of SCZ-altered and APD influenced proteome. **A)** A representative image of western blot analysis for *GLS* proteins associated with the SCZ-altered proteome in an independent cohort. Valosin-containing protein (*VCP*) was used as loading control. **B)** Significant (*p-value* < 0.02) down-regulation of *GLS* in the MS-based analysis. **C)** Western blot analysis of *GLS* in an independent cohort did not show any significant difference. However, the downregulation was significant (*p-value* < 0.03) for subjects within 20 to 37.5 years of age. **E)** Hypergeometric overlap between the SCZ-altered and APD influenced proteome with similar proteome studies of the DLPFC (top), other brain regions (middle) and drug specific features (bottom). The size of circles is proportional to the −log10 of p-values associated with the overlap. The black dotted circles represent the reference = −log10(*p-value* = 0.05).

Overall, the differential expression analysis reveals a conserved set of protein regulated across several brain areas and studies; however, expression of these proteins in present and other studies is dependent on drug effect.

### Distinct functional changes associated with SCZ and APD

Next, to understand the functional changes associated with SCZ-altered and APD-influenced proteome, we performed gene-ontology (GO) analysis (aka: functional pathway analysis, Figure 3). Identified pathways (*q-value* < 0.05) were clustered into functional themes (Figure 3, left labels). Upregulated pathways were involved in only a few themes related to *organelle*, *vesicles*, and *extracellular region*. The *extracellular region* was the most prominently altered upregulated theme, as the majority of other pathways (Supplementary Table 4) including those involving *organelles* and *vesicles* (*extracellular exosome*, *extracellular organelle*, *extracellular vesicle* and *neurofilament*) also have extracellular functionalities. Notably, the majority of upregulated pathways overlapped with those that were downregulated. Given that GO pathways represent a set of coordinated features^35–37^ against different biological processes, the simultaneous enrichment of extracellular functionalities in the up- and down-regulated feature-sets reflects their aberrant coordination in SCZ.

**Figure 3:**
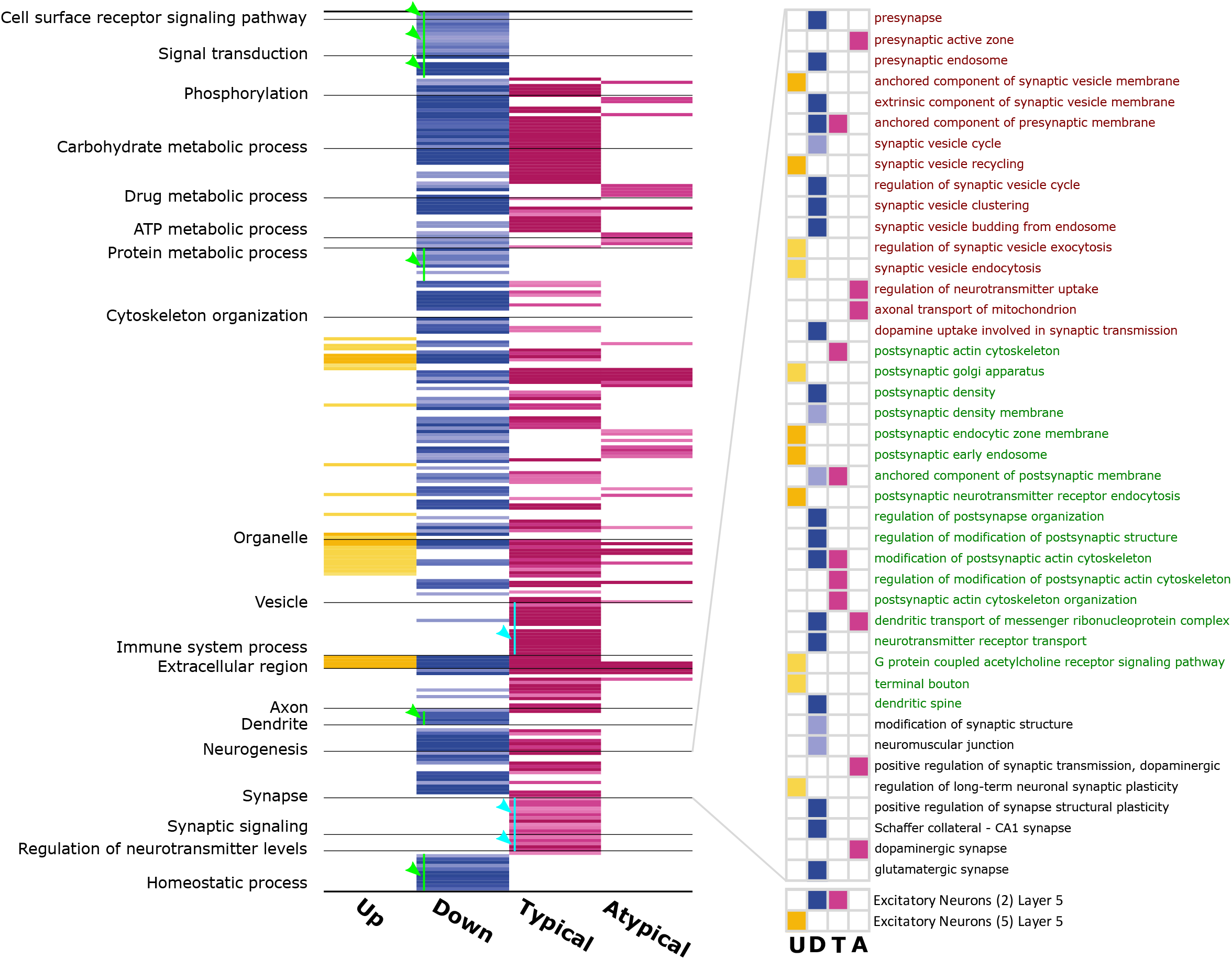
Biological pathways, synaptic changes and cell-types associated with SCZ-altered and APD influenced proteome. Changes associated with SCZ-altered up- and downregulated proteomes shown in yellow and blue, respectively. Pathways associated with typical and atypical APD influenced proteome are shown in pink. The green arrowheads indicate pathways exclusive to the SCZ-altered downregulated proteome, whereas the cyan arrowheads indicate pathways specific to typical APDs. Left inset (top): The presynaptic (red), postsynaptic (green) and other synaptic pathways (black) studied using SynGo database anchoring curated synapse related ontology. Left inset (bottom): cell-type specific enrichment of SCZ-altered and APD influenced proteome studied using DLPFC specific cell-type signatures. Lighter to darker shade of any color represents increasing enrichment score represented by −log10(*q-value*). See Supplementary Table 4 for details.

Downregulated pathways were involved in almost all the themes and overlapped largely with typical APD associated pathway; however, there were some notable non-overlaps between the two profiles. For instance, the downregulated pathways showed exclusive enrichment of themes associated with *cell surface receptor signaling*, *signal transduction*, *dendrites*, *cytoskeleton organization* and *homeostatic* process (Figure 3, green arrowhead); whereas the typical APD associated pathways showed exclusive enrichment of themes associated with *synaptic signaling*, *regulation of neurotransmitter levels* and *immune system process* (Figure 3, cyan arrowhead). Lastly, the atypical APDs, showed fewer pathways and were mostly associated with themes involving *extracellular region*, *vesicles*, *organelle*, *phosphorylation*, and metabolic process involving ATP, drug, and carbohydrate metabolism. Unlike the typical APDs, atypical APDs showed no association with immune system process and other structural changes (*cytoskeleton organization*, *axons*, *dendrites and synaptic signaling*).

Overall, the functional analysis reveals that, with the exception of the loss of homeostasis, signal transduction, dendritic, and cytoskeleton-related processes, APDs appear to compensate for the majority of the pathways disrupted in SCZ but alter synaptic signaling and the immunological process.

### Distinct synaptic events and layer 5 pyramidal neurons are disrupted in SCZ

Disruptions of synaptic and cellular function plays an important role in the complex network of events that underpins SCZ-pathophysiology^38^. To zoom-in the enrichment of those events in the functional analysis, we used SynGO^24^, a database of synaptic ontology (Figure 3, inset top)and cell-specific signatures from human DLPFC^25^ (Figure 3, inset bottom). Synaptic enrichments (*p-value* < 0.05) were organized into presynaptic (Figure 4, red) and postsynaptic (Figure 4, green) groups of pathways.

**Figure 4:**
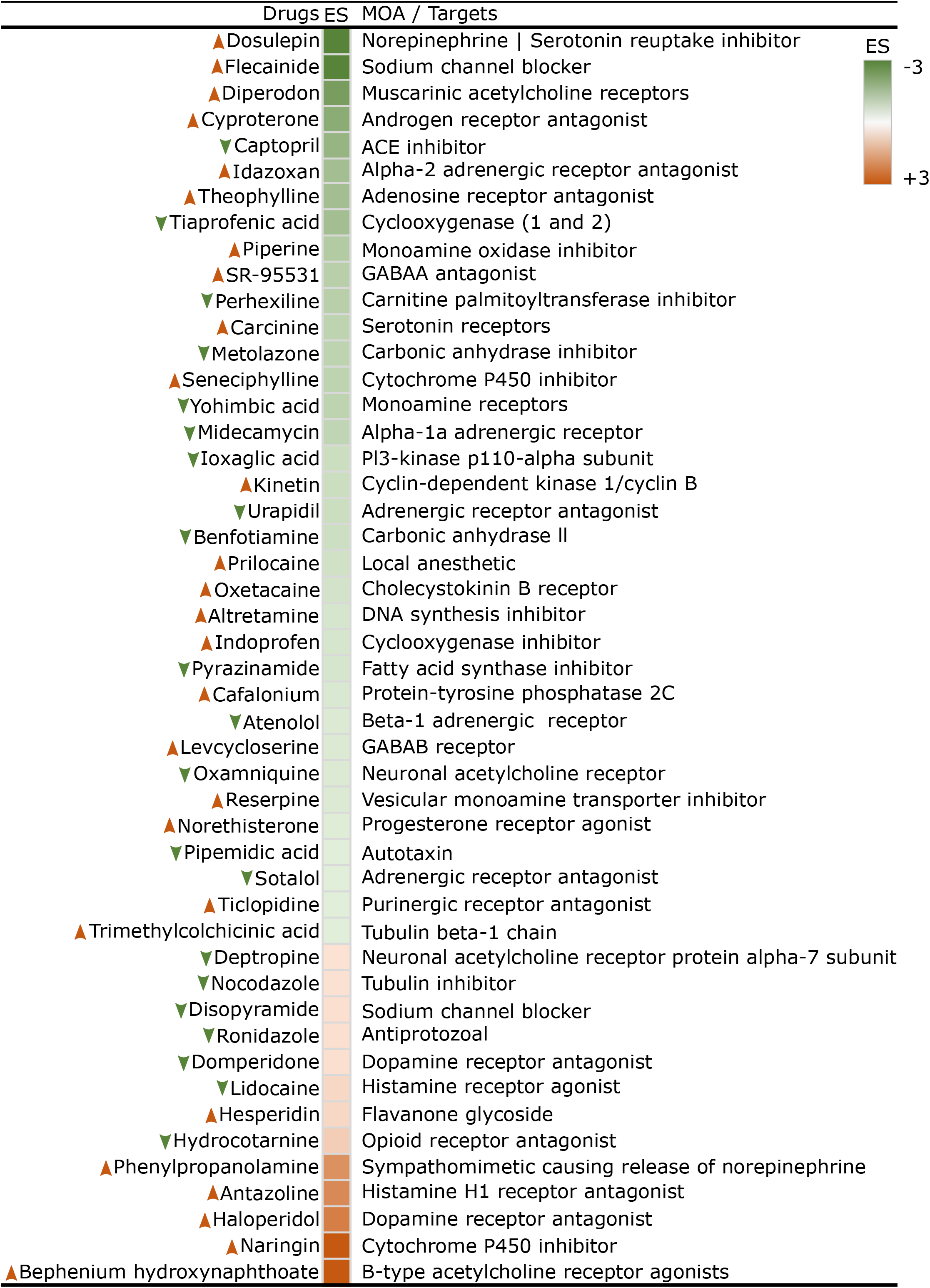
Drug and MOA/target associated with SCZ-altered proteome. The orange and green boxes represent the enrichment (ES) of drugs with known MOA/targets in up- and down-regulated proteomes, respectively. A lighter to darker shade of both orange and green represents an increasing enrichment calculate using −log10(*p-value of enrichment statistics*). The upward-pointing orange arrow and downward-pointing green arrow represent the up- and down-regulated drug signatures, respectively, as predicted by the cmap database. See Supplementary Table 7 for details.

Downregulated proteins were equally enriched in presynaptic and postsynaptic components. Postsynaptic components, however, exhibited greater structural variability (*actin-cytoskeleton*, *mRNA complexes*, *neurotransmitter receptor transport and anchored and membrane components of post synaptic density*) as opposed to presynaptic components, which were primarily associated with the synaptic vesicle and its regulation. Besides pre- and postsynaptic changes, *dopamine-uptake* and *glutamatergic-synapse* associated pathways, were also downregulated.

Upregulated proteins, on the other hand, were more enriched in the postsynaptic components associated with *Golgi apparatus*, *endocytic zone*, *endosomes*, and *G-coupled acetylcholine receptors*. Upregulated presynaptic components were associated with vesicle movements (*exocytosis*, *endocytosis*, and *recycling*). Long term synaptic plasticity was also upregulated. Among the synaptic changes associated with different APD signatures, atypical APD were mostly associated with presynaptic components including dopaminergic synapse and neurotransmitter uptake. Mitochondrion-related pathways were also found in conjunction with atypical APDs, implying that these medicines impact bioenergetic functions. Typical APDs in contrast were primarily associated with postsynaptic components linked to cytoskeleton and its organization.

Within the different cell-types specific signatures obtained from DLPFC specific single-cell studies^25^, both up- and downregulated proteins were enriched in a non-overlapping set of layer-5 pyramidal neurons (PNs). Characterizing these neurons further (see Supplementary Table 6 notes) using GO revealed that the PN subsets were exclusively associated with regulation of dopamine (Excitatory-Neurons-(2)-Layer-5: *q-value* < 5.02 × 10^−5^; Excitatory-Neurons-(5)-Layer-5: *q-value* < 9.12 × 10^−4^). Additionally, based on enrichment of different deep layer projection neuron markers^39^, these neurons appear to be striatum projecting (Excitatory Neurons (2) Layer 5: *p-value* < 0.08; Excitatory Neurons (5) Layer 5: *p-value* < 0.09).

Overall, zooming in on cellular and synaptic changes reveals a balance of pre- and postsynaptic changes, as well as the influence of up- and down-regulated proteins on distinct subsets of dopamine-regulated striatal projecting layer-5 PNs associated with distinct neuromodulatory and synaptic events.

### Insights into the key MOA associated with SCZ specific signatures

To understand the molecular events (MOA or targets) that precede the aforementioned functional events, we looked for enrichment of drug-signatures associated with known MOAs/targets in SCZ-altered up- and down-regulated proteins. 62 and 18 drugs were enriched (*p-values* < 0.05) in up- and down-regulated proteins, respectively (Figure 4, Supplementary Table 7 notes).

MOAs/targets involving upregulated events includes neuromodulation (*dopamine*, *norepinephrine*, and *acetylcholine*), local immune response (*histamine*), and regulation of pain (*opioid receptors*). Contrary to the signature reversing principle^40^, which assumes discordance between drug-disease signatures for therapeutic effect, several known SCZ-drugs were concordant with disease signatures, confirming that SCZ-altered proteome, as noted, are likely to be driven by the medications.

Among the MOAs/targets shared by up- and down-regulated proteins, several neuromodulatory events (except dopaminergic), hyperpolarization events (*sodium channel blockers*), and cytochrome p450 activity were observed, implying that these targets are critical pharmacological nodes in SCZ pathophysiology. MOAs exclusive to down-regulated events include: both fast (GABA-A) and slow (GABA-B) inhibitory modulations; serotonergic and norepinephrine modulation by means of receptors and transporters; different cell-surface and cytoplasmic signal transduction events (*tyrosine-protein kinase LCK*, *Serine/threonine-protein kinase*); enzymes involving homeostasis (*carbonic anhydrases*), fatty acid oxidation (*carnitine palmitoyl transferase*), and anti-inflammatory activity (*cyclooxygenase*); hormonal receptor activity involving gender specificity (*androgen* and *progesterone*); cellular processes inhibiting the depolarization by means of *acetylcholinesterase*; cellular processes associated with DNA metabolic process (*DNA alkylation, DNA synthesis*); microtubule organization (*tubulin beta-1 chain*) and cellular detoxification events (*Glutathione S-transferase kappa 1*) and gastrointestinal regulation (cholecystokinin B receptor).

Overall, SCZ-altered upregulated events were linked with a few MOA/targets, most of which were neuromodulatory, whereas downregulated events were linked with a more diverse range of MOA/targets, outside from neuromodulation.

## Discussion

The majority of SCZ subjects receive APD treatment, which limits the inferences drawn from postmortem examinations. Here, to understand the effect of SCZ and APDs (in SCZ), we contrasted the statistically derived SCZ-altered and APD-influenced proteome using functional analysis focusing on GO, cell-types, and subcellular synaptic changes. Using drug-specific signatures, we demonstrate that the majority of SCZ-altered changes were indeed influenced by APDs. However, our approach could mitigate this issue in two different ways. First, the contrast between pathways associated with SCZ-altered and APD-influenced proteome revealed that homeostasis, signal transduction, cytoskeleton, and dendrite related processes were not much compensated for by APDs during SCZ. The latter two processes (cytoskeleton and dendrite) are consistent with the increased neuronal density in non-treated postmortem SCZ-DLPFC, which is accompanied by a decrease in cell size (attributable to cytoskeleton) and dendritic spines^41, 42^. Second, signatures of drug with known MOA revealed potential mechanism involved in SZC pathophysiology despite the confounding effect of drugs. For instance, besides the influence of drug in our cohort, an upregulated therapeutic effect of dopamine receptor antagonist subscribes to the known compensatory upregulation of dopamine receptors during SCZ pathophysiology^43^. Interestingly, the SCZ altered proteome in the present study significantly overlapped with several previous SCZ related findings across different studies and brain region. Furthermore, functional analysis using these proteomes demonstrate several key SCZ specific finding, for instance, effect of extracellular region^44^, association of layer-5 PNs with SCZ pathology^45^ and its projection to striatum^46^ which are consistent with previous several molecular, anatomical, and functional imaging-based studies. While these consistencies validate our approach and reproducibility of data, the influence of APDs in our study calls in to question the inference made based on these prior studies.

### Influence of SCZ-altered and APD-influenced proteome in cortico-striatal pathways

DLPFC neurons (Figure 5) project to *dorsal striatum* related to associative functionality via cholinergic interneurons which modulates dopaminergic input (to striatum) through the nicotine receptors presents in it^47^. The output of *dorsal striatum* includes the direct and indirect projections to basal ganglia (*globus pallidus internal*) via the GABAergic medium spiny neurons (MSN)^48^. MSNs projecting directly uses D1 (excitatory) receptors and those relaying indirectly via *globus pallidus external* and *subthalamic nucleus* uses D2 (inhibitory) receptors. The *globus pallidus internal* projections hyperpolarize the thalamus which projects back to layer-3 where the inhabitant PNs extend axon collaterals to other cortical areas^43^. Additionally, DLPFC, similar to dorsal striatum, receives input form the dopaminergic neurons of ventral mesencephalon^49^.

**Figure 5:**
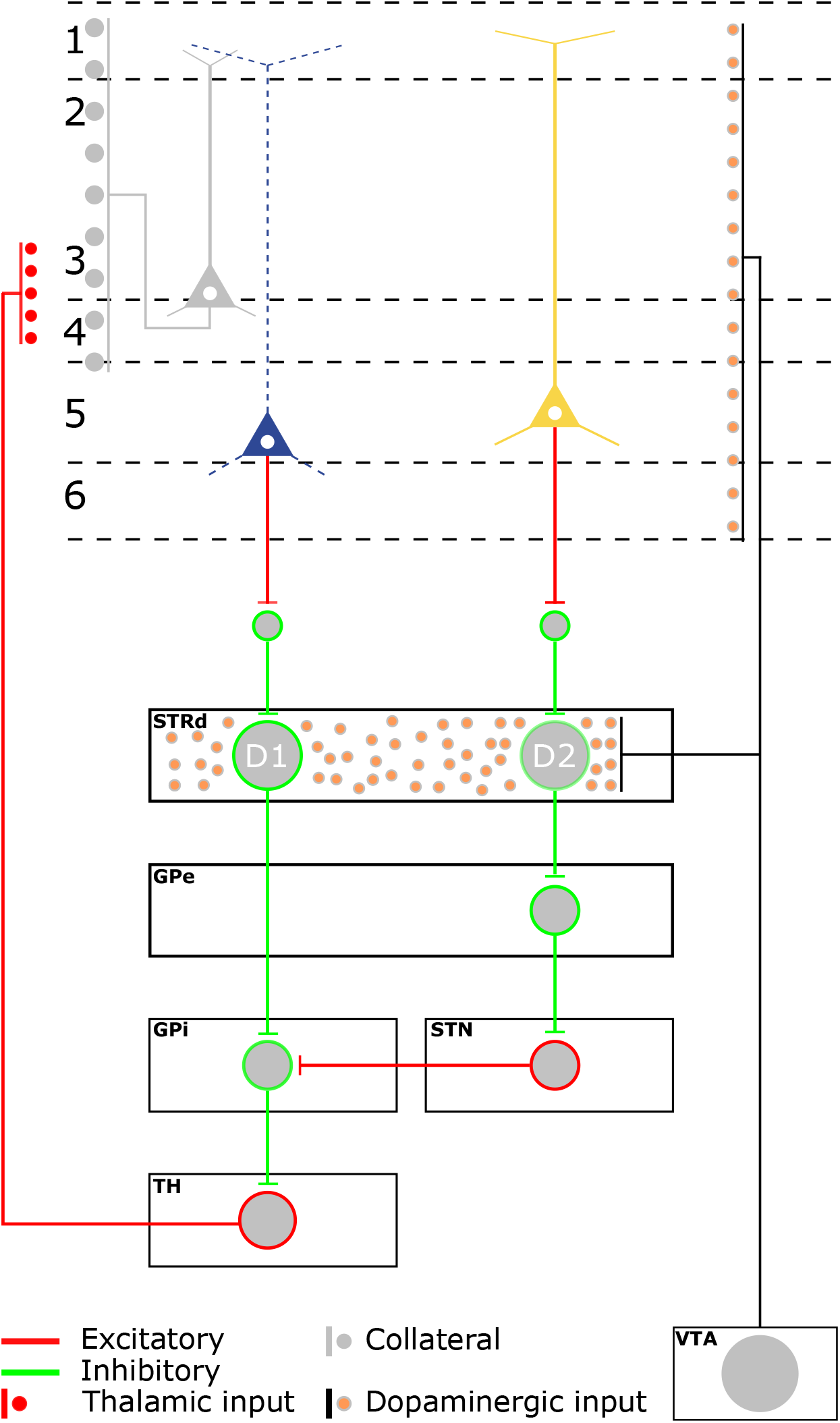
Schematic summary of functional changes observed in the study in the context of cortico-stratum circuitry. Up- and down-regulated SCZ-altered proteome were enriched in two different subset of Layer 5 PNs showing enriched dopaminergic signaling. The PNs enriched in upregulated events were also associated with D2 receptor antagonist. Together these point towards selective association of the SCZ-altered up- and downregulated proteome with D1-PN and D2 PN of cortical layer 5, respectively. The data consistent with previous anatomical studies also suggests that the two neurons are striatum projecting, perhaps to the D1 and D2 expressing MSN neurons. An order of excitatory and inhibitory events shown in red and green, respectively sum up to modulate the excitatory thalamic input to the cortex. Unlike previous studies pustulating a hypodopaminergic cortex the functional analysis also suggests a hyperdopaminergic cortex and a potential selective and compensatory pruning of D1-PN dendrites (shown as a dashed blue line). STRd: dorsal Striatum, GPe: Globus pallidus external, GPi: Globus pallidus internal, STN: Subthalamic nucleus, TH: Thalamus, VTA: Ventral tegmental area. See text for more details.

Our functional analysis reveals that the up- and down regulated changes are enriched in two different subsets of striatum projecting layer 5 PNs, which based on the exclusive enrichment of dopaminergic signaling can be linked to D1- and D2-receptor expressing cortical PNs. PN with upregulated event (Figure 5, yellow) based on the association with *haloperidol*, which targets D2 receptor^50^ can be the D2-PN while the PN with downregulated events (Figure 5, blue) can be the D1-PN. In normal circumstances when dopamine is optimal, the network events sum up to relay usual thalamic input to cortex (Figure 5, legend). However, during SCZ, the hyperactivity of dopamine in striatum assigns salience to unremarkable environmental stimuli^51^ and relays it to cortical layer-3 PNs^52^ (Figure 5, grey PN) whose collaterals could cascade the salience signal to other cortical areas leading to an inappropriate affect. Note that within this mechanism (i.e., hyperdopaminergic striatum), previous studies have postulated a hypodopaminergic cortex^43, 53–56^; but our functional analysis based on downregulated dopamine uptake (Figure 3) and upregulated dopamine receptor antagonists (Figure 4) suggests that the hyperdopaminergic effect leading to SCZ-pathology might simultaneously initiate in the cortex and the striatum. A possible and perhaps important pathological implication of this could be: first, a maladaptive association between working memory (a DLPFC function^57^) and salience (a striatum function^58^), which can be explored in SCZ patients using functional MRI. Note that in normal conditions, this association can be between working memory and reward^59^. Second, an association between stress— a prefrontal cortex (PFC) related functionality and SCZ. Supporting the latter, a recent study on mouse chronic unpredictable stress demonstrated that D1-and D2-PN subpopulations of PFC undergo distinct stress-induced intrinsic and synaptic plasticity changes that may have functional implications for stress-related pathology^60^. One can argue that a hyperdopaminergic cortex, being consistent with most medication-based mechanism (e.g., *Haloperidol*), is likely relevant to the positive and negative symptoms of the SCZ (inappropriate affect) but does not enumerate the cognitive deficit related to DLPFC functionality. This, we conjecture, is largely due to the fact that cognition (e.g., working memory) may be associated with the intracellular signaling events^61, 62^ while most SCZ-relevant therapeutic changes are targeting the receptors events implicated in a circuit wide phenomenon. We observed several downregulated kinase related events in our MOA/target analysis which are involved in memory formation^63^. In this regard a more rational design might look for adjuvants which targets the kinase pathways suggested here (Figure 4).

Further placing the functional analysis results in the context of a hyperdopaminergic cortex, the reduced dendrite and cell-size (cytoskeleton, Figure 3) seems to be selective to the D1-PNs (blue) and not to D2-PNs or other PNs (particularly layer-3) as postulated before^43^. As the postulated hyperdopaminergic cortex by means of D1-PNs may also sum up to relay inappropriate affect from the thalamus, the selective pruning of these D1-PN-dendrites and reduced cell size may be a compensatory mechanism which is further supported by the equally distributed pre- and post-event— a phenomenon often associated with the compensatory response^43^. Lastly, although, our study focused only on DLPFC, we observed conserved result across several brain area suggesting that a similar hyperdopaminergic mechanism might appears in all areas; however, depending upon the location can have different functionality. For instance, in the auditory or motor cortex and related area in striatum the inappropriate affect instead of amplifying the salience can amplify inner speech leading to thought echo a phenomenon consistently observed in SCZ patient^48^.

### Limitations

First, besides the fact that several variable including sex, age, and race plays an important role in SCZ pathophysiology^64^, the data was not stratified or regressed for these variables as this would have reduced the power of the analysis. Studying the effect of these variable along with drug-effect individually can be perused in future. Second, the functional predictions obtained through bioinformatics analyses should be interpreted as hypothesis-generating.

## Author contributions

R.S. and R.S.A. conceptualized the study and together wrote the manuscript. J.R, J.M and R.E.M participated with R.S.A in LCMS data generation and processing. S.M.O and A.J.F participated with R.S.A in western blot analysis. All authors participated in writing the manuscript.

## Data and materials availability

All analyzed data are available in the main text or the supplementary materials. Raw data are available from the corresponding author on reasonable request.

## CONFLICT OF INTEREST

The authors declare no competing interests.

Supplementary information is available at MP’s website.

