## Supplementary Information for "Pharmacological impacts on schizophrenia functional analysis: a postmortem proteome study"

### Supplementary Materials and methods

#### Protease digestion and electrospray tandem mass spectrometry

Twenty micrograms of protein for all samples were solubilized in Laemmli sample buffer (6X solution: 375 mM Tris-HCl, 9% SDS, 9% beta mercaptoethanol, 50% glycerol and 0.03% bromophenol Blue, pH 6.8, Fisher Scientific, Massachusetts) then heated at 60 ºC for 10 minutes. Each sample was run approximately 2 cm into a preparative mini 1D gel, excised and digested with trypsin as previously described[1]. The extracted peptides were analyzed by nano liquid chromatography coupled to electrospray tandem mass spectrometry (nLC-ESI-MS/MS) performed on a 5600+ QTOF mass spectrometer (Sciex, Toronto, On, Canada) with an Eksigent (Dublin, CA) nanoLC ultra nanoflow system, as used previously [2], but with the addition that samples were prepared in such that 2.5 ug of protein digest containing 1X of iRT (indexed Retention Time) and internal standard (#Ki-3002-1;Biognosys, Boston, USA) was used for each injection.

#### Data- dependent Acquisition (DDA) mode

The DDA system was operated in positive ion mode with collection over the 90 min gradient run. Each cycle included 1 time of flight (TOF) mass spectrometry scan covering the m/z 400-1250 window with a 200 ms accumulation time followed by isolation and fragmentation of the 50 most intense ions with charge states of +2 to +4 with an accumulation time of 40 ms resulting in a full duty cycle of just under 2.25 sec. Former MS/MS-analyzed candidate ions were excluded for 12 s after their first occurrence to reduce the redundancy of the identified peptides.

#### Data- independent Acquisition (DIA) mode

For the DIA method, we used SWATH-MS as prescribed previously[2], where each 2.5 ug sample was injected and separated using the same 90 min gradient as the DDA. Then subjected to a custom series of overlapping mass windows for data collection that cover a range of m/z between 400-1250, including a 1 Da overlap of successive windows (Table S3). This produced

1704 cycles where each duty cycle included 1 TOF-MS scan covering m/z 400-1250 at 200 ms accumulation time followed by 100 DIA segments each with a 28 ms accumulation time for a total duty cycle of 3.05s. The NanoSpray III ion source (Sciex) parameters used for all analyses included a source gas 1 (GS1), source gas 2 (GS2), and curtain gas (CUR) set to 8, 0, 35, respectively. The system was operated in positive ion mode with the heated interface and the ion spray voltage set to 150C and 2.6kV, respectively. MS/MS spectra were collected using the high sensitivity mode with the rolling collision energy function with an energy spread of 5V as provided in the Analyst-TF v. 1.7 (Sciex) data acquisition software.

**Western blot analysis**

Western blot was used to examine the expression of malate dehydrogenase 1 (*MDH1*), glutaminase (*GLS*). Ten micrograms of protein samples were prepared in Laemmli sample buffer at 70 ºC for 10 min. Samples were run in duplicate into 17-well pre-cast 4-12% Bis-Tris gel (NuPAGE Invitrogen, ThermoFisher Scientific) in 1X MES buffer (Invitrogen, ThermoFisher Scientific) for 1 hour at 180 V. The following gel was transferred to a polyvinylidene fluoride membrane for semidry transfer (Bio-Rad) for 30 minutes at 18 V current using a semi-dry transfer apparatus (Bio-Rad). The membranes were blocked for 1 h at room temperature with Licor blocking buffer in 1X tris-buffered saline with tween 20 (TBS-T). All membranes were incubated with primary antibody in blocking buffer with 0.2% Tween20 either mouse anti-MDH1 (1:500, AM31915PU-N, OriGENE), rabbit anti-glutaminase (1:5000, ab156876, Abcam), or a housekeeping control rabbit anti- valosin containing protein (*VCP*,1:1000, ab109240, Abcam) at 4 ºC for overnight. Membranes were washed three times in TBST with 0.01% Tween20 for 10 min before being probed with anti-mouse (1:5000, #68070, Licor) or anti-rabbit (1:5000, #68073, Licor) IR-dye labeled secondary antibodies in blocking buffer with 0.2% Tween 0.01% SDS for 1 h at room temperature in the dark. Membranes were then washed again with TBST for three times each for 10 min then scanned by using the LI-COR Odyssey laser-based imaging system. Image Studio 4.0 software used to measure each band intensity values with segment median intra-lane background subtraction. Following that, each band was normalized to the calibrator sample (pool of all sample) present in each blot as an internal control and normalized to *VCP* as reference protein. We pre-tested the *MDH1*, *GLS*, and *VCP* using varying concentrations of protein from a human tissue homogenate sample.

**Supplementary table (each as sperate excel file) legends and notes**:

**Supplementary Table 1: Subject demographics.**

- Sheet [Cohort 1 (MS)]: Cohort used for mass-spectrometry based proteomics analysis.
- Sheet [Cohort 2 (WB)]: Cohort used for wester-blot based validation.

For both cohorts brain samples were obtained from Maryland Brain Collection.

**Supplementary Table 2: SCZ-Altered proteome.** Differentially expressed proteome identified using moderate t-statistics using the limma package in R. Details of the column header is provided below:

- **MS ID:** Protein identification in MS run
- **Proteins:** Protein symbols. The non-redundant differentially expressed list of up and down regulated proteins were used for all functional analysis described in the text
- **LogFC:** log fold change between CLT and SCZ subjects
- **AveExpr:** Average MS intensity across the CLT and SCZ samples
- **t:** t-statistics
- **P.Value:** the non-adjusted p-value used for filtering the SCZ-altered proteome.
- **Adj.P.Val:** FDR corrected p-value (due to fewer proteins passing the threshold of 0.05, this was not used for filtering the SCZ-altered proteome.

**Supplementary Table 3: Proteomes influenced by the demographics.** Output of *variancePartition* package in R[3]. The values represent the per protein (rows) variance explained (in decimals), for each available covariate (columns). The violin plot shown in Figure 1A was generated using data for selected covariates with the highest average variance. For typical and atypical APDs proteins showing > 10 percent variance (shown in red) were selected for all downstream functional analysis.

**Supplementary Table 4: Functional analysis using Gene Ontology (GO, sheet 1) and SynGO (sheet 2).** List of GO and SynGO (synaptic ontology) terms (rows) associated with

SCZ-altered up- and down-regulated proteomes as-well-as typical and atypical APD-influenced proteome (columns). For GO all pathways with *q-value* < 0.05 are shown. For SynGO all pathways with *p-value* < 0.05 are shown. The values represent *enrichment score (ES)* = -*log10(q/p-value)*. For both GO and SynGO all pathways were clustered into themes. The table was used to generate Figure 3.

**Supplementary Table 5: Gene sets that were used to conduct customized functional analysis.**

- **Sheet [Similar Studies]:** Gene-sets were curated from similar studies focusing SCZ associated proteomes from DLPFC[4, 5] and different brain areas[6-11].
- **Sheet [PN markers]:** gene markers associated with deep layer PNs[12-23] projecting to different brain areas. These makers were used to characterize the DLPFC neuronal makers used from *Nagy et.al.*[24] See notes for Supplementary Table 6.

For both the gene sets, enrichment analysis was performed as described in material and methods sections using gene-overlap package in R. For gene-set associated with *Similar Studies*, only significantly enriched gene-sets were used to plot Figure 2F (top and middle).

**Supplementary Table 6: Characterization of PNs.** We examined the enrichment of DLPFC cell-specific signatures available from Nagy et al. in SCZ-altered and APD-influenced proteomes. Two distinct subsets of PNs were discovered to be enriched. We ran two further enrichment analyses to further characterize them. First, we analyzed all of the DLPFC cell-specific signatures using GO analysis (**Sheet: Functional characterization**). Notably, only the two groups of PNs enriched in the SCZ-altered proteome (Ex_2_L5 and Ex_5_L5, highlighted in orange) were exclusively enriched in dopaminergic signaling (pathway highlighted in orange), indicating that these neurons might be the layer-5 D1 and D2 expressing PNs. Second, to determine the area to which these PNs project (**Sheet: Anatomical characterization**), we used the signatures of PNs (Supplementary Table 5, sheet: PN markers) projecting from the cortex to other brain regions. The two PNs were enriched in signatures associated with striatum projecting layer-5 PNs with trend level significance (relevant p-value, highlighted in green).

**Supplementary Table 7: List of Drugs with known MOA/Targets enriched in up- and down regulated SCZ-altered proteome.** Details of the column header is provided below:

- **Direction:** The up (orange) and down (green) arrows indicate whether the drug signature enriched in the SCZ-altered proteome is upregulated or downregulated, respectively. Note that signature form comparative toxicogenomics database (ctd) do not have direction. All drugs signature with directions are form connectivity map (cmap) database.
- **Drugs:** Name of the drug whose signatures are significantly enriched in the SCZ-altered proteome.
- **Enrichment Score (ES):** -log10(*p-value* < *0.05* of enrichment score). The green boxes (negative ES) represent enrichment of drug signature in down regulated SCZ-altered proteome while the orange boxes (positive ES) represent enrichment of drug signature in up regulated SCZ-altered proteome.
- **MOA/Target:** Mode of action / targets associated with each drug molecule.
- **Notes:** Details regarding the target molecule gathered from various internet resources

Note that based on signature reversion principal an up- and down-regulated drug matching with the respective up- and down-regulated disease signature (i.e., green arrow matching green box and orange arrow matching orange box) suggests a drug with disease mimicking effect while up- and down-regulated drug matching with the down- and up-regulated disease signature (i.e., green arrow matching orange box and orange arrow matching green box) suggests a drug with disease antagonizing effect. However, *Haloperidol*, a disease antagonist, was identified as a disease mimicking drug, implying that the SCZ-altered proteome is not independent of drug effect.  Only drugs with directionality are shown in Figure 4.
